# Cell type-specific effects of isoflurane on two distinct afferent inputs to cortical layer 1

**DOI:** 10.1101/2020.05.18.102913

**Authors:** Caitlin A. Murphy, Matthew I. Banks

**Affiliations:** Department of Anesthesiology, University of Wisconsin, Madison, Wisconsin, United States

**Keywords:** isoflurane, anaesthesia, inhalation, thalamus, neocortex, interneurons, consciousness, patch-clamp techniques, channelrhodopsin

## Abstract

**Background:** While their behavioral effects are well-characterized, the mechanisms by which anaesthetics induce loss of consciousness are largely unknown. Anaesthetics may disrupt integration and propagation of information in corticothalamic networks. Recent studies have shown that isoflurane diminishes synaptic responses of thalamocortical (TC) and corticocortical (CC) afferents in a pathway-specific manner. However, whether the synaptic effects of isoflurane observed in extracellular recordings persist at the cellular level has yet to be explored.

**Methods:** Here, we activate TC and CC layer 1 inputs in non-primary mouse neocortex in *ex vivo* brain slices and explore the degree to which isoflurane modulates synaptic responses in pyramidal cells and in two inhibitory cell populations, somatostatin-positive (SOM+) and parvalbumin-positive (PV+) interneurons.

**Results:** We show that the effects of isoflurane on synaptic responses and intrinsic properties of these cells varies among cell type and by cortical layer. Layer 1 inputs to L4 pyramidal cells were suppressed by isoflurane at both TC and CC synapses, while those to L2/3 pyramidal cells and PV+ interneurons were not. TC inputs to SOM+ cells were rarely observed at all, while CC inputs to SOM+ interneurons were robustly suppressed by isoflurane.

**Conclusions:** These results suggest a mechanism by which isoflurane disrupts integration and propagation of thalamocortical and intracortical signals.

## INTRODUCTION

Anaesthetics may ultimately influence both the level and contents of consciousness via actions on corticothalamic circuits, disrupting integration of information throughout the cortical hierarchy^1-3^. Activity in higher order cortical areas is particularly sensitive to anaesthetics^4, 5^, as is corticocortical (CC) feedback connectivity^6-11^. In addition to cortical effects, higher order thalamocortical (TC) connectivity is suppressed during both sleep^12^ and anaesthesia^13-15^. Activation of non-specific thalamic nuclei during anaesthetic-induced unconsciousness^16^ and non-REM sleep^17^ promotes behavioral arousal, implicating higher order thalamic centers as enablers of consciousness.

Recent studies from our lab have shown that in *ex vivo* slices, isoflurane diminishes synaptic responses in a pathway-specific manner^10, 11^. In the latter study, despite overlapping terminal fields in layer 1, CC feedback afferents to a non-primary cortical area were preferentially suppressed by isoflurane compared to non-primary TC inputs. However, extracellular recordings used in these investigations capture summed effects across all synapses, leaving open questions as to whether observed effects are consistent across post-synaptic targets. For example, differential effects on inhibitory interneurons compared to excitatory cells may shift the balance of excitation and inhibition^18, 19^, disrupting integration of inputs in pyramidal cell dendrites or constraining the output of the cortical column. Furthermore, superficial and deep layers may operate within functionally separate domains, where cells in superficial layers primarily carry out intracortical interactions, and those in deep layers facilitate interactions with lower cortical and subcortical targets in the feedback direction^20-23^. Differential effects of anaesthetics on cells residing in different layers may provide insight into the functional roles of discrete cortical networks and the mechanisms that subserve the breakdown of connectivity observed during anaesthesia^24^.

Here, we investigate the effect of isoflurane on independently activated TC and CC inputs to layer 1 of non-primary neocortex in brain slices. We evaluate the pathway- and cell type-specific effects of isoflurane on synaptic responses recorded in pyramidal cells in layers 2-5 and in two nonoverlapping subpopulations of inhibitory cells, SOM+ and PV+ interneurons.

## METHODS

### Selective activation of afferent pathways to non-primary visual cortex

To express the fluorescent reporter tdTomato in non-overlapping populations of GABAergic interneurons, homozygous, Cre-dependent tdTomato male mice were bred with either SOM-Cre female or PV-Cre female mice; heterozygous offspring (both sexes) were used for all experiments. Viral constructs containing channelrhodopsin-2 and fluorescent reporter YFP (AAV2-hSyn-hChR2(H134R)-eYFP; provided by K. Deisseroth, Stanford University via University of North Carolina Vector Core) were injected into either posterior medial thalamus (AP: -2.25 mm, ML: 1.25 mm, DV: -3.35 mm; volume: 1 µL) or cingulate cortex (AP: 0.2 mm, ML: 0.3 mm, DV: 0.9 mm; volume: 1 µL) to activate independently non-primary TC and CC feedback afferents, respectively.

After allowing at least 3 weeks for expression of the virus, acute coronal brain slices containing were collected and recordings were made from medial secondary visual cortex. To activate either TC or CC afferents, blue light was delivered exclusively to layer 1 (150 µm diameter circle, 470 nm; Polygon400, Mightex Systems, Toronto, Ontario). For each trial, synaptic responses were evoked using four brief pulses of light (2 msec; 10 Hz) at one of at least five light intensities randomly selected from a range of intensities (0.022-2.2 mW). Each trial was separated by ten seconds, and at least eight trials were conducted for each light intensity, randomly interleaved.

### Electrophysiological recordings and data analysis

Intracellular whole cell patch clamp recordings were conducted in current clamp mode from pyramidal cells in layers 2-5, SOM+ interneurons, or PV+ interneurons; extracellular multichannel recordings were collected concurrently. Cell types were identified either by Cre-dependent expression of td-Tomato reporter (for SOM+ and PV+ interneurons) or by morphology (pyramidal cells). To further guide identification of cell types, each cell’s response to 600 msec current steps was evaluated as previously described^25^ (Supplementary Table 1). All signals were low-pass filtered at 10 kHz and digitized at 40 kHz. For each cell, recordings were conducted for control, isoflurane (0.24-0.29 mM), and recovery conditions, with 20 minutes between drug conditions to allow for equilibration.

Intracellular signals were bandpass filtered (0.1-1000 Hz), and responses were averaged across trials of the same stimulus intensity. Signals from each of 16 extracellular channels were bandpass filtered (1-300Hz) to yield local field potentials, from which the current source density^26^ (CSD; spline inverse method^27^) was calculated. The CSD signal in layer 1 with the shortest latency current sink (inward transmembrane current) was used for further analysis.

For both intracellular and extracellular recordings, responses to a single stimulus intensity were selected for detailed analysis; the intensity selected was the minimum light intensity for which at least 50% of trials (8-10 trials per intensity) evoked a response with <1 msec latency jitter. Response amplitude was defined as the peak of the signal within 50 msec of the stimulus (first pulse only). Paired pulse ratio (PPR) was calculated by dividing the amplitude of the response to the second pulse by the amplitude of the response to the first pulse. TC- and CC-evoked responses were compared across control, isoflurane (0.24-0.29 mM), and recovery conditions (Figures 1-3). To exclude the possibility that effects of isoflurane were due to response degradation over time, the average of control and recovery measures were used as a baseline for comparison to isoflurane in the statistical model.

**Figure 1.**
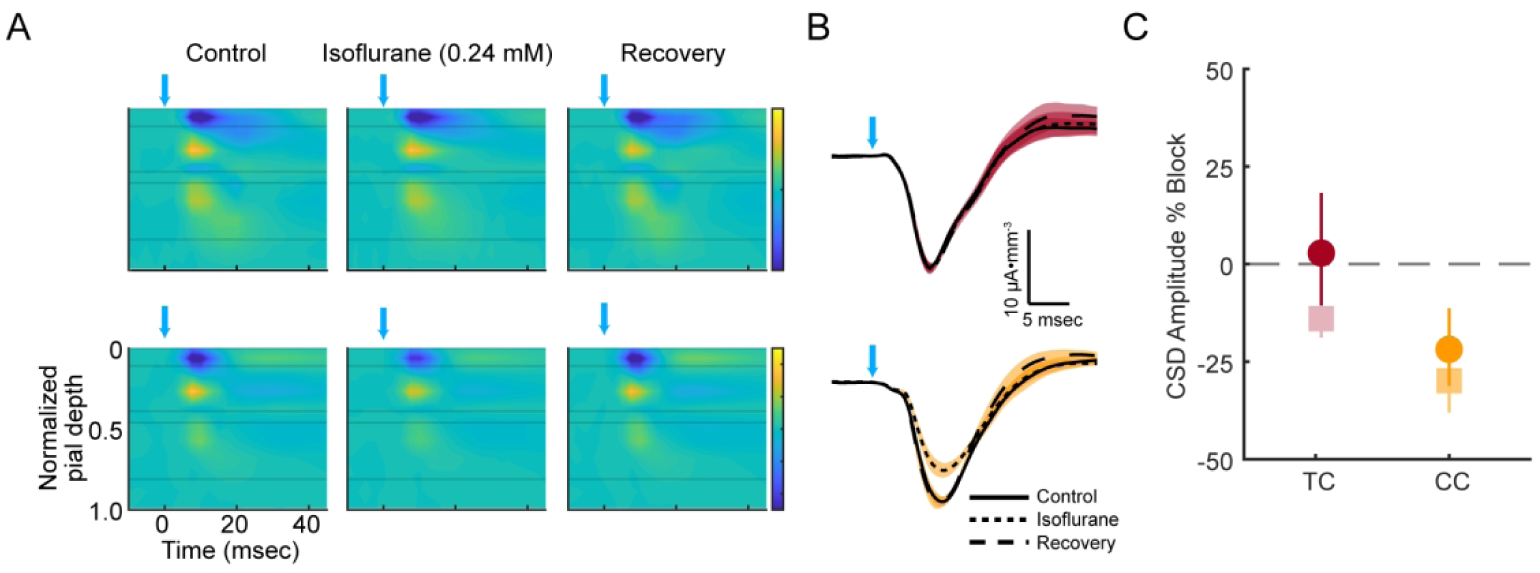
Effects of isoflurane on optogenetically-evoked current sinks in layer 1 for TC and CC inputs. *A*. CSD color plots comparing evoked synaptic responses to a 2 msec light pulse (blue arrows) across control (*A*, left column), 0.24 mM isoflurane (*A*, middle column), and recovery (*A*, right column) for representative examples for TC (top row) and CC (bottom row) afferents; color bar: -12.5 to 12.5 µA mm ^-3¬^, where blue current sinks indicate inward-going transmembrane current. *B*. Signals from layer 1 were used to compare synaptic effects across control (solid), 0.24 mM isoflurane (dotted), and recovery (dashed) for TC (top) and CC (bottom) inputs. Shaded regions indicate ± 1 SD among trials. *C*. Model fits (see Methods) of percent block of layer 1 sink by isoflurane with 95% confidence interval (vertical line). Responses evoked by stimulation of CC afferents to cortical layer 1 (orange) are suppressed by isoflurane, while TC inputs to layer 1 (red) are not. Pale square markers reflect estimates of block by isoflurane at 0.24 mM based on results from Murphy et al., 2019. Raw values of CSD sink amplitudes under control conditions are reflected in **Table 1**.

### Statistical analysis

A linear mixed effects model was used to evaluate the pathway- and cell type-dependent effects of isoflurane on intrinsic and evoked response properties of individual cells. Fixed effects were cell type (L2/3 Pyr, L4 Pyr, L5 Pyr, SOM+, or PV+) and drug condition (baseline or isoflurane), as well as afferent pathway (TC or CC) for evoked responses. Interaction terms between isoflurane and each of the other fixed effects were also included, as well as a three-way interaction term representing interactions between cell type and pathway of drug effects; random effects were slice experiment and cell with random slopes for drug condition. Likelihood ratio tests are reported using chi-squared values. The significance of individual coefficients was tested and isoflurane effects are reported relative to baseline with 95% confidence intervals. For amplitudes of post-synaptic potentials (PSPs) and input resistance measures, heteroscedasticity was corrected by log transforming response variables, and effects are reported as percent changes.

**Table 1.**
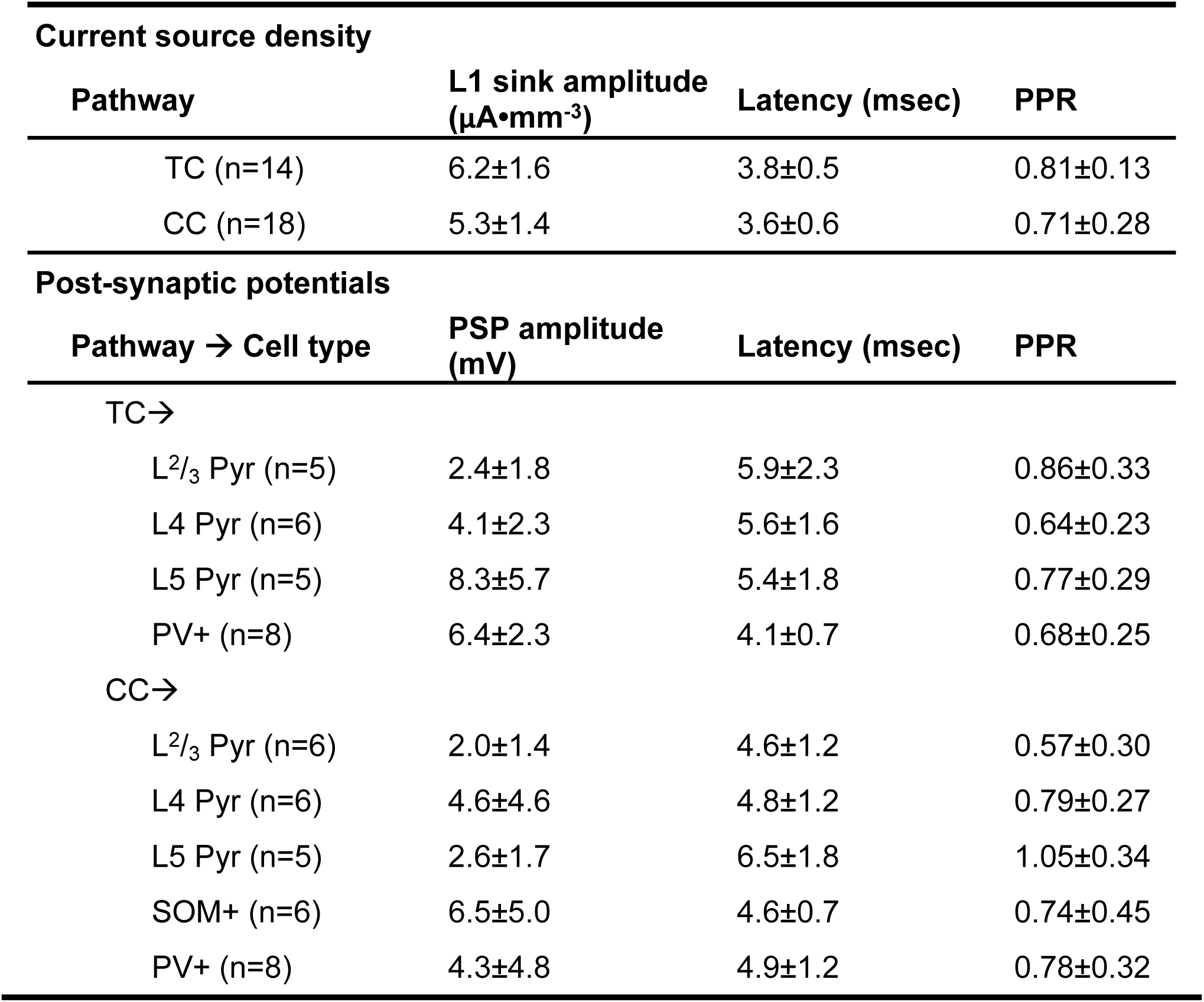
Properties of evoked synaptic responses during control conditions. Optogenetic activation of TC and CC terminals evoked current sinks in L1 (top) and post-synaptic potentials in non-overlapping cell populations (bottom). Values shown are mean ± standard deviation. Latency was calculated relative to onset of 2 msec light pulse. Paired pulse ratios were calculated as the amplitude of the 2^nd^ response (relative to the pre-response baseline) divided by the baseline-subtracted amplitude of the 1^st^ response.

## RESULTS

### Activation of TC and CC inputs to layer 1 elicit short-latency synaptic responses

Synaptic responses from TC and CC afferents were elicited by activating channelrhodopsin in afferent axon terminals in layer 1. Brief pulses of light evoked short latency current sinks in layer 1 (Table 1). No effect of afferent pathway was observed for baseline amplitude (t(29) = 0.41, p=0.69) or latency (t(29) = 0.26, P = 0.80) of layer 1 current sinks. Evoked currents in layer 1 precipitated PSPs in pyramidal cells in layers 2-5 and in SOM+ and PV+ interneurons (Table 1).

Main effects of pathway and cell type on amplitudes of synaptic responses were tested using averaged responses from the lowest stimulus intensity need to elicit a response. A 2-way ANOVA was not significant for effects of pathway (F(1,47) = 0.07, P=0.79) nor target cell population (F(4,47) = 1.68, P=0.17), suggesting no differences in baseline (i.e., in the absence of isoflurane) amplitudes among recorded cells. Similarly, no significant effect on latency was observed for main effects of pathway (F(1,47) = 1.00, P=0.32) nor cell type (F(4,47) =1.68, P=0.17).

### Isoflurane suppresses layer 1 current sinks elicited by CC inputs, but not TC inputs

Despite their overlapping terminal fields and similar synaptic dynamics under control conditions, TC and CC inputs to layer 1 were differentially affected by isoflurane (Figure 1; Table 2), consistent with previous results from our lab^11^. Amplitudes of layer 1 current sinks were suppressed by isoflurane by 21.8% for CC inputs, while current sinks evoked by TC inputs were relatively resistant to isoflurane (Figure 1; Table 2).

**Table 2.**
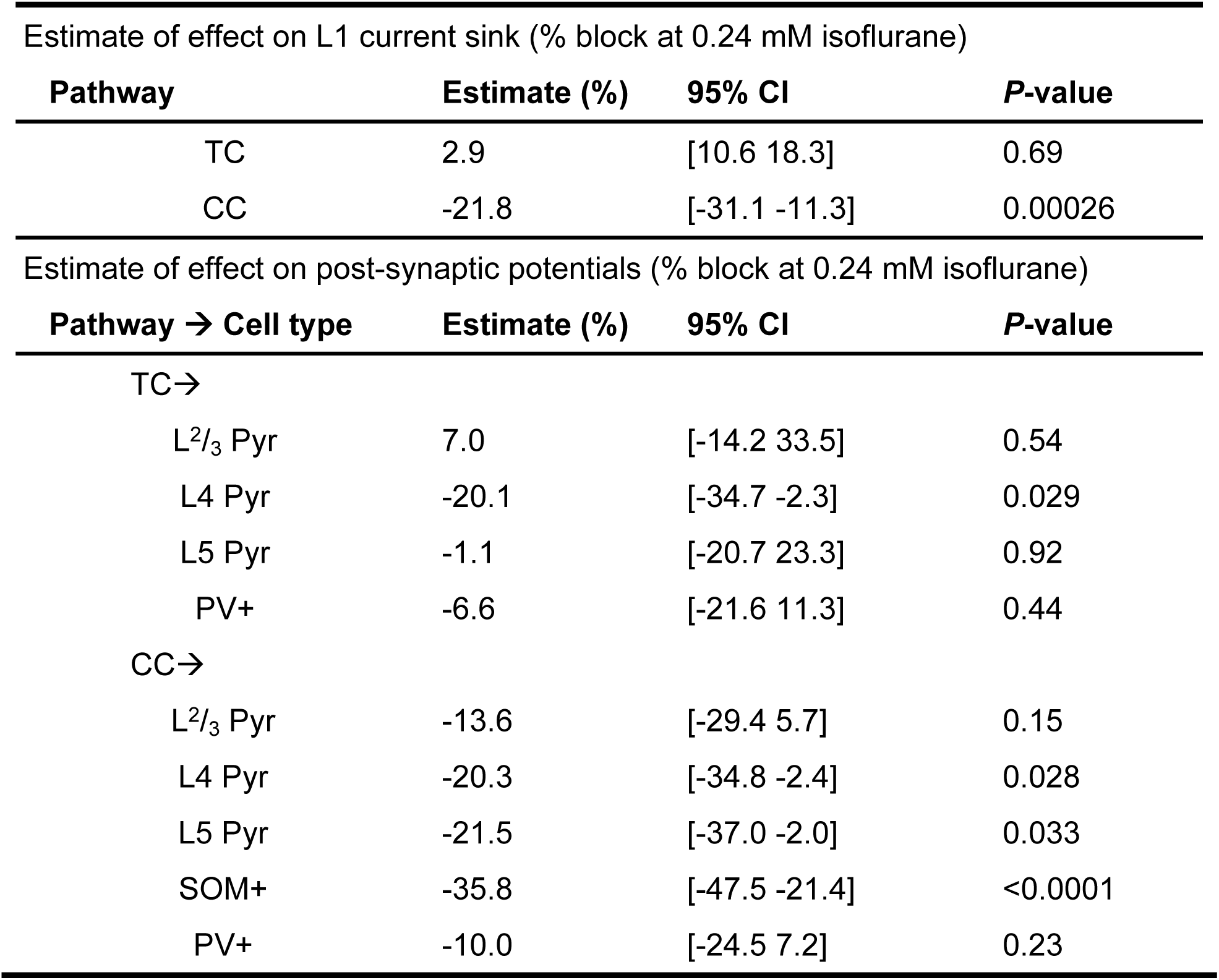
Isoflurane affects synaptic responses in a pathway- and cell type-dependent manner. Percent block by isoflurane (0,24 mM) of layer 1 current sink (top, from extracellular CSD signals) and post-synaptic potentials (bottom, from intracellular current clamp signals) evoked by activation of TC and CC inputs. Estimates are based on predicted values from linear mixed effects model (see Methods). Values represent the percent change in the amplitude of current sinks or post-synaptic responses during isoflurane relative to baseline conditions. Negative values represent suppression of synaptic responses by isoflurane.

### Isoflurane suppresses synaptic responses in a cell type-dependent manner

Current sinks isolated from current source densities represent averages of inward-going transmembrane currents across all synapses. However, evidence suggests that the synaptic properties – and therefore potential functional roles – of TC versus CC inputs to may be cell type- and layer-dependent^28, 29^. Therefore, we sought to investigate whether the effects of isoflurane on layer 1 current sinks were uniform for all post-synaptic targets of each afferent pathway, or whether certain cell populations were differentially sensitive to isoflurane.

We evaluated the degree to which isoflurane modulated the amplitudes of PSPs elicited by layer 1 afferents in L^2^/_3_, L4, and L5 pyramidal cells (Figure 2) as well as SOM+ and PV+ cells (Figure 3). Independent of any interactions of cell type or afferent pathway, isoflurane significantly suppressed evoked post-synaptic potentials of target cells (χ(1) = 16.6, P<0.0001). Contrary to our findings for extracellularly recorded layer 1 current sinks, we found no significant interaction of pathway on the effect of isoflurane (χ(1) = 2.00, P=0.16). However, the effect of isoflurane was significantly dependent on cell type (χ(4) = 17.862, P=0.0013). The interaction between cell type and pathway was not significant (χ(1) = 6.52, P=0.16). In other words, the effect of isoflurane is cell type-dependent, yet within each cell type, no significant differences were detected between afferent pathways.

**Figure 2.**
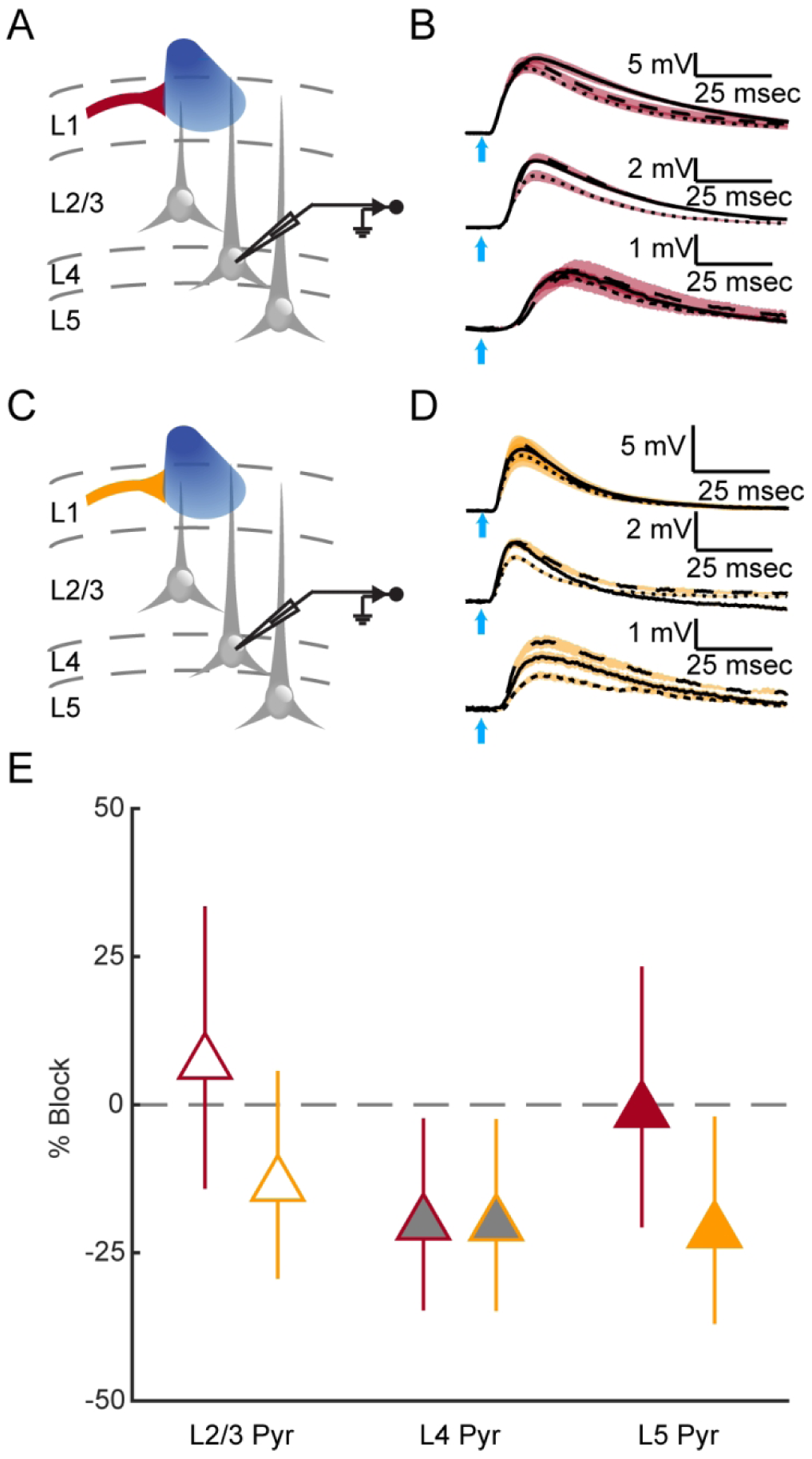
Example recordings from cortical pyramidal cells in response to activation of two distinct layer 1 afferents. *A*. Schematic demonstrating recordings conducted for panel B. *B*. Example recordings of TC-evoked responses from pyramidal cells in L2/3 (top), L4 (middle), and L5 (bottom) during control (solid), 0.24 mM isoflurane (dotted), and recovery (dashed) conditions. Shaded regions indicate ± 1 SD among trials. *C-D*. Same as *A-B* for responses to CC inputs. Raw values for control conditions are shown in **Table 1**. *E*. Linear mixed effects model estimates (see Methods) of the percent block by isoflurane of post-synaptic responses evoked by TC (red) or CC (orange) inputs to L2/3, L4, and L5 pyramidal cells. Values represent the percent change in the amplitude of post-synaptic responses during isoflurane relative to baseline conditions; negative values represent suppression of synaptic responses by isoflurane. Vertical lines span 95% confidence intervals for isoflurane effects for each pathway-cell type combination. Both TC and CC inputs to L4 pyramidal cells were suppressed by isoflurane (TC→L4 cells: -20.1% [-34.7, -2.3], P=0.029; CC→L4 cells: -20.3% [-34.8, -2.4], P=0.028), as were CC inputs to L5 pyramidal cells (CC→L5 cells: -21.5% [-37.0, -2.0], P=0.033). P-values for all pathway-cell type combinations are shown in **Table 2**.

**Figure 3.**
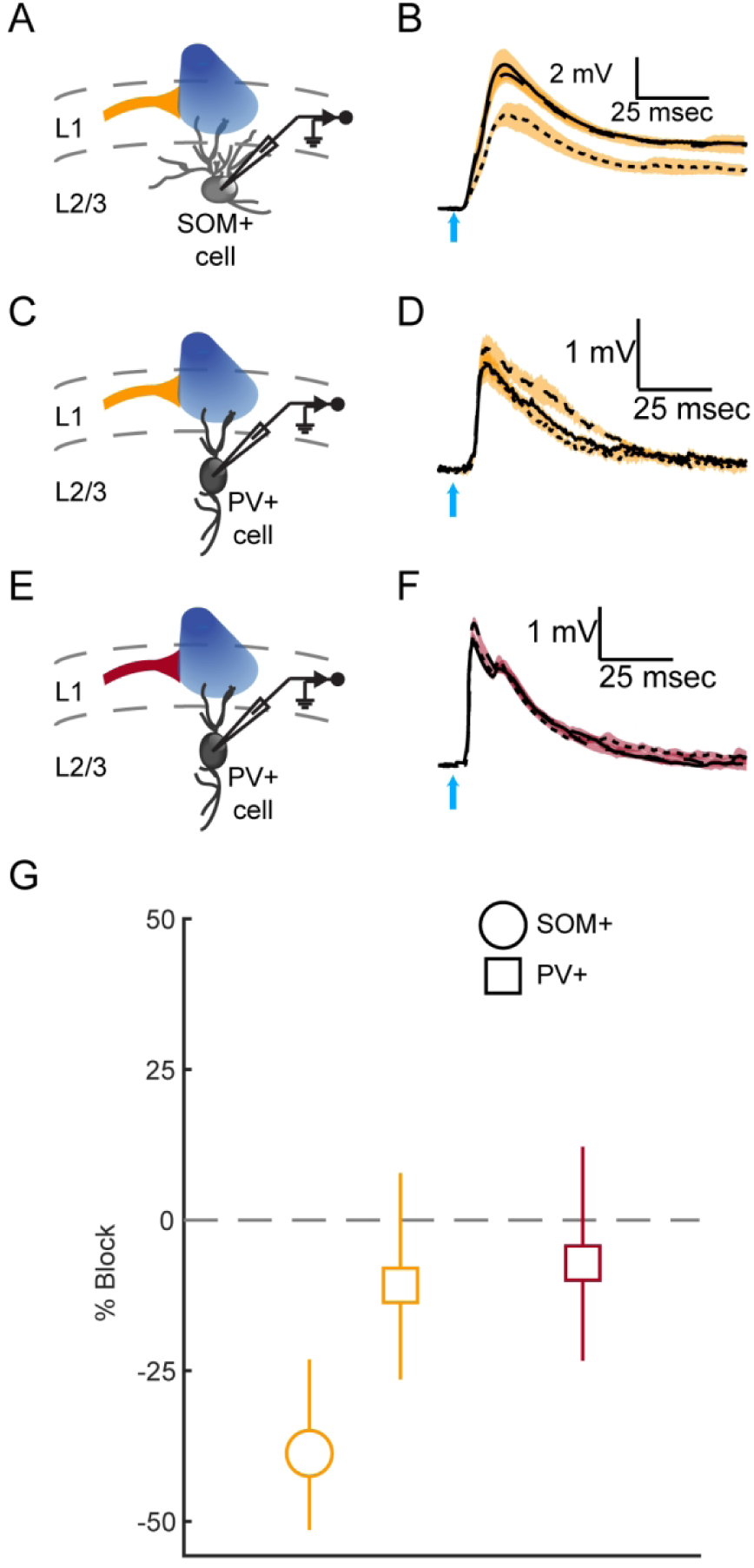
Isoflurane suppresses inputs to SOM+ and PV+ interneurons in a cell type-dependent manner. *A*. Schematic demonstrating recordings conducted for panel B. CC afferents to layer 1 were optogenetically activated while whole cell patch clamp recordings were conducted in SOM+ interneurons in L2/3. *B*. Example recordings from CC-evoked responses in a SOM+ cell during control (solid), 0.24 mM isoflurane (dotted), and recovery (dashed) conditions. Shaded regions indicate ± 1 SD among trials. *C-D*. Same as *A-B* for CC inputs to PV+ cells. *E-F*. Same as *A-B* for TC inputs to PV+ cells. Note that examples of TC inputs to SOM+ cells are not shown, as TC inputs failed to reliably evoke synaptic responses in SOM+ cells. Raw values of PSP amplitude during control conditions are shown in **Table 1**. *G*. Linear mixed effects model estimates of the percent block by isoflurane of post-synaptic responses evoked by TC (red) or CC (orange) inputs to SOM+ (circle) and PV+ (square) interneuron populations. Vertical lines span 95% confidence intervals for isoflurane effects for each pathway-cell type combination. CC inputs to SOM+ cells were suppressed by isoflurane (CC→SOM+ cells: -35.8% [-47.5, -21.4], p<0.0001), but no effect of isoflurane was detected in TC nor CC inputs to PV+ cells. P-values for all pathway-cell type combinations are shown in **Table 2**.

Although there was a difference in the effect of isoflurane on responses to TC versus CC inputs in L2/3 pyramidal cells (Figure 2; Table 2), neither effect was significantly different from zero. Conversely, amplitudes of synaptic responses in L4 cells were significantly suppressed by isoflurane for both TC and CC inputs, to approximately the same degree (Figure 2; Table 2). Isoflurane also suppressed CC inputs, but not TC inputs, to L5 pyramidal cells, though this effect was not significantly different between pathways (P=0.19).

In addition to providing input to pyramidal cells, long-range TC and CC projections also target inhibitory interneurons. We were interested in the effect of isoflurane on evoked responses in two subpopulations of inhibitory cells, SOM+ and PV+ interneurons, and whether these effects were different than those observed for pyramidal cell populations. Consistent with previous studies^30-32^, TC inputs rarely elicited synaptic responses in SOM+ cells (n=2 cells); as such, isoflurane effects on TC inputs to SOM+ cells are not reported. Activation of CC afferents, however, reliably elicited synaptic responses in SOM+ cells (Figure 3). These synapses were particularly sensitive to isoflurane (Figure 3); amplitudes of synaptic responses to CC inputs in SOM+ cells were suppressed 35.8% during isoflurane compared to baseline (Table 2). Conversely, no effect of isoflurane on amplitudes of responses in PV+ cells was detected for either afferent pathway (Figure 3; Table 2). Thus, even among subpopulations of GABAergic interneurons, the effect of isoflurane on CC responses was significantly different (−35.8% for SOM+ cells versus -10.0% for PV+ cells, P=0.014; Figure 3), suggesting distinct roles of each cell type during anaesthesia.

### Pre- and post-synaptic effects of isoflurane are cell type-dependent

A multitude of cellular and synaptic dynamics may underlie changes in neural activity observed during anaesthesia. For example, effects of isoflurane on synaptic responses as we describe here may result from pre-synaptic effects of isoflurane (e.g., effects on calcium influx or vesicle release^33^), from effects on discrete intrinsic properties of post-synaptic cells that give rise to changes in their excitability, or a combination of pre- and post-synaptic effects. Although we cannot ascribe the contributions of specific synaptic or intrinsic effects of isoflurane to observed effects on evoked responses, characterizing isoflurane-induced changes in short-term synaptic plasticity or intrinsic properties of post-synaptic cells may provide insight into the complicated interactions of network components during anaesthesia. To this end, we evaluated the extent to which isoflurane modulated PPR (Figure 4) and intrinsic membrane properties (Supplementary Figure 1) of each cell relative to baseline. Baseline values are reflected in Table 1 for short-term plasticity and in Supplementary Table 1 for intrinsic parameters.

**Figure 4.**
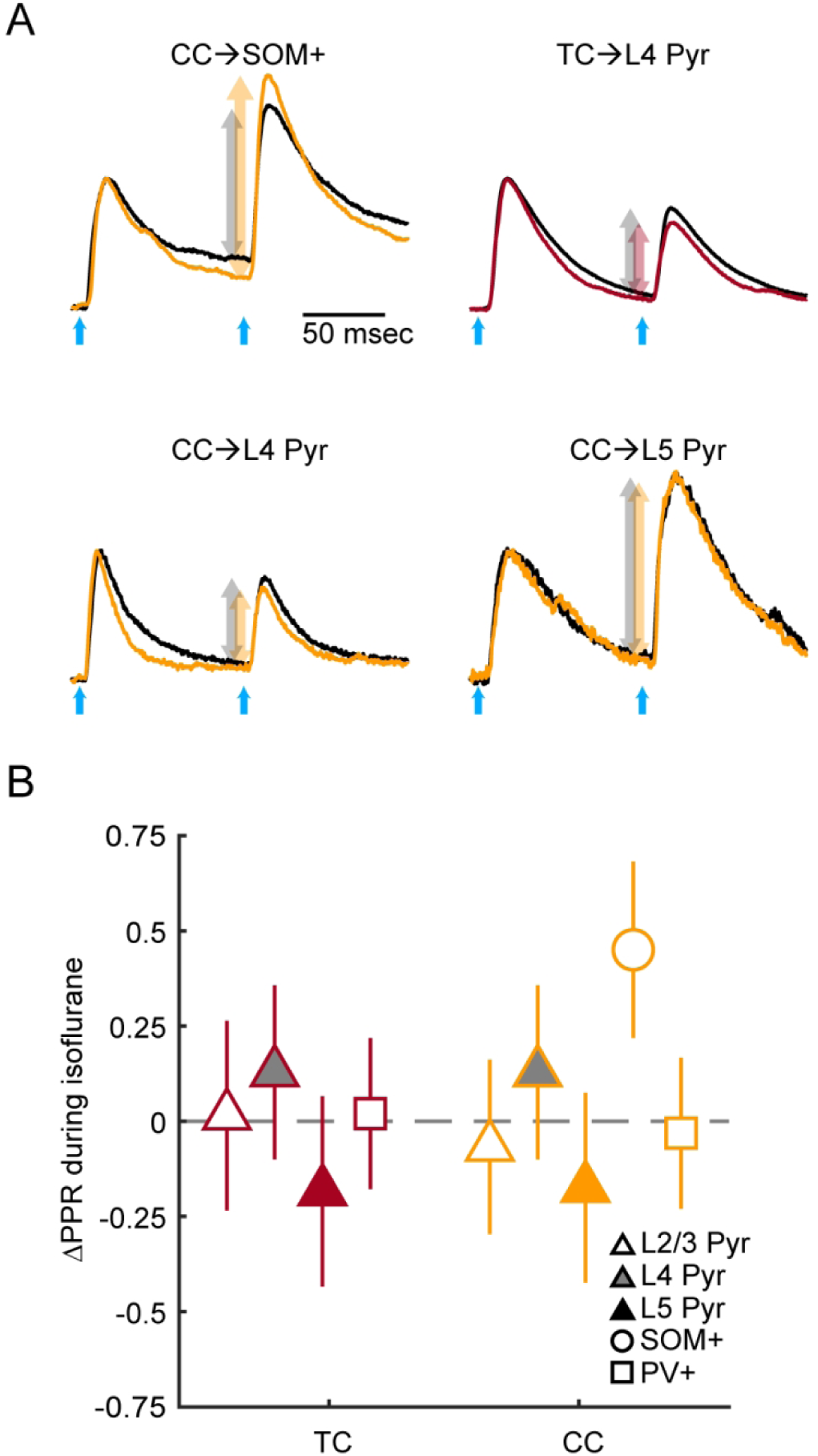
Effect of isoflurane on short-term plasticity for TC and CC inputs to pyramidal cells and interneurons. *A*. Example recordings from the four different pathway-cell type pairs that demonstrated suppression of synaptic responses by isoflurane. Responses for control (black) and isoflurane (either orange (CC) or red (TC)) conditions are normalized to the amplitude of the first pulse under control conditions to demonstrate paired pulse effects. Arrows compare paired pulse effects between control (black) and isoflurane (either orange or red) conditions. *B*. Model estimates of the changes to PPR during isoflurane, where positive values represent larger PPR during isoflurane compared to baseline. Estimates of isoflurane effects on PPR for responses evoked by TC (red) or CC (orange) inputs to all recorded populations are shown. Vertical lines span 95% confidence intervals for isoflurane effects for each pathway-cell type combination. PPR for CC inputs to SOM+ cells were enhanced by isoflurane (CC→SOM+ cells: +0.45 [+0.22, +0.68], p=0.0002), suggesting isoflurane blocks synaptic responses to SOM+ cells through pre-synaptic mechanisms. Raw values of PPR during control conditions are shown in **Supplementary Table 2**.

Isoflurane significantly enhanced the PPR of CC inputs to SOM+ cells but had no effect on PPR for other pathway-cell type combinations (Figure 4; Supplementary Table 2). Intrinsic membrane properties of L4 pyramidal cells were especially sensitive to isoflurane (Supplementary Figure 1). In L4 pyramidal cells, input resistance was 13.5% lower (95% CI [-23.4 -2.4], P=0.019) and membrane time constants 27.8% shorter (95% CI [-42.3 -9.5], P=0.0051) during isoflurane relative to baseline; isoflurane did not affect input resistance nor time constants of any other cell type (Supplementary Figure 1).

## DISCUSSION

In these experiments, we describe the effects of isoflurane on synaptic responses and intrinsic membrane properties of pyramidal cells in layers 2-5, as well as in SOM+ and PV+ interneurons, in non-primary visual cortex. Numerous studies have shown that feedback cortical connectivity is preferentially suppressed by anaesthetics^6-11^. Results from extracellular recordings we present here recapitulate these findings: we show that extracellular synaptic responses to activation of layer 1 inputs are pathway-dependent, with CC feedback responses suppressed and TC responses unaffected. While our extracellular results here corroborate canonical paradigms, our observations at the cellular level tell a different story. We demonstrate that the suppression of synaptic responses by isoflurane is cell type- and layer-dependent, where responses to both TC and CC inputs to layer 4 pyramidal cells are suppressed and those to PV+ inhibitory interneurons are unaffected.

Despite TC and CC feedforward *connectivity* being relatively preserved during anaesthesia, *activity* in higher areas – the targets of higher order TC and CC feedforward connections – is suppressed^4, 5, 34^. Could cell type-specific effects of anaesthetics explain this paradox? As is suggested by the discrepancy between our own extracellular and intracellular results, extracellular recordings may not be robust enough capture specific effects at the cellular level. Understanding effects of anaesthetics on discrete circuit components not only provides insight into mesocircuit-level effects underlying the mechanisms of anaesthesia, but may also provide structure for understanding the enigmatic relationship between “activity” and “connectivity”. Further examination of the implications of cell type-specific effects may allow reconciliation of previous conclusions drawn from investigations using seemingly disparate methods.

### Resistance of PV+ interneurons to isoflurane suggests shift in E/I balance

We show that synaptic responses of PV+ interneurons to both TC and CC inputs are resistant to suppression during isoflurane. TC input to PV+ cells provides stimulus-locked fast feedforward-inhibition to pyramidal cells, restricting the window of opportunity for generating spikes and improving spike timing among populations of pyramidal cells^25, 35^. We propose that the relative insensitivity of fast-spiking PV+ neurons to isoflurane we observe here may underlie previous observations from our lab and others in which early responses of the cortical network to TC inputs are relatively preserved during isoflurane, while recurrent, propagating network activity is highly sensitive^24, 36, 37^. Although we report activity of single cells only in our study, the relative preservation of PV+ cell responses during isoflurane is likely to shift the E/I balance in the network toward inhibition. PV+ cells are known to be important for mediating E/I balance within cortical networks^38, 39^. Pyramidal cells in L2/3, for example, receive PV+ cell-mediated inhibitory input proportional to their excitation under normal conditions, and changes in input to pyramidal cells – as we observe in L4 cells in our experiments – without commensurate changes in PV+ cells disrupts this E/I balance^18^. If these cell type-specific effects of isoflurane occur in multiple cortical areas, shifts in local E/I balance may compound, contributing to the progressive failure of information to be passed from one cortical area to the next.

### Suppression of L4 pyramidal cells during isoflurane may disrupt feedforward processing

We show that synaptic responses of L4 pyramidal cells are suppressed during isoflurane for responses evoked by both TC and CC inputs. That TC inputs to any cell type would be affected by isoflurane is somewhat surprising, and that this effect would be exclusively observed in L4 pyramidal cells is even more unexpected. L4 pyramidal cells are typically considered mediators for propagating feedforward information, receiving sensory-specific information from thalamus and lower areas^31, 40-43^ and targeting L2/3 for output to higher areas^44, 45^. Feedforward circuits are relatively resistant to anaesthetics^10, 11, 46^ (though, see ^47, 48^), and we show here that extracellular recordings of current sinks evoked by activation of TC afferents were not affected by isoflurane (Figure 1). Yet, interestingly, our results show that both TC and CC inputs to L4 cells are preferentially suppressed by isoflurane (Figure 2), suggesting that suppression of either afferent pathway may disrupt feedforward signaling.

While pre-synaptic mechanisms of isoflurane have been well-described^33, 49, 50^, our observations suggest that the mechanisms underlying isoflurane effects in L4 pyramidal cells were not limited to pre-synaptic effects. No changes in short-term synaptic plasticity were observed at these synapses (Figure 4), and TC and CC inputs to L4 pyramidal cells were suppressed to a similar extent (Figure 2). However, L4 pyramidal cells were significantly leakier during isoflurane, as evidenced by lower input resistances and shorter membrane time constants relative to baseline (Supplementary Figure 1). Not only might these effects dampen voltage changes evoked by synaptic currents, but they are likely to also contribute to a decrease in the overall excitability of L4 cells, thereby attenuating their contributions to processing of cortical signals.

### Evidence for disruption of predictive coding schemes during anaesthesia

In the present study, we find that isoflurane suppresses CC feedback afferents to L4 and L5 pyramidal cells (Figure 2) as well as those to dendrite-targeting SOM+ interneurons (Figure 3), consistent with previous studies showing sensitivity of feedback connections during anaesthesia^6, 7, 10, 11, 46^. Feedback activation of the cortical column is mediated largely by intracortical afferents to layer 1 that target distal dendrites of pyramidal cells. Dendritic calcium currents activated by these inputs increase the gain of the input/output relationship of cortical pyramidal cells and are highly correlated with sensory perception^51, 52^. This process is tightly regulated by powerful disynaptic inhibition from SOM+ interneurons^53^. Our observation that inputs to both SOM+ cells and infragranular pyramidal cells are preferentially suppressed notably intersects with these previous findings in the context of sensory coding. Suppression of intracortical feedback inputs to two key players in this integration process may therefore disrupt the typical input/output function of pyramidal cells.

Disruption of integration of inputs to L5 pyramidal cells is also consistent with what might be expected under anaesthesia in the context of current theoretical models. The predictive coding hypothesis, for example, has often been used as a framework for understanding consciousness and sensory processing^54-56^. Under the predictive coding hypothesis, feedback signals must activate inhibition to offset incoming excitatory signals and prevent propagation of the excitatory, ascending error signal. Discrepancies between internally generated predictions and sensory input from external sources are propagated as excitatory “error signals”. Our data provide further evidence that multiple components of the predictive coding architecture may be disrupted by anaesthesia. First, integration of predictions carried by feedback afferents may be perturbed, as feedback inputs to both SOM+ cells and L5 pyramidal cells – key mediators of feedback modulation of sensory responses – are altered by isoflurane. Second, propagation of error signals in the feedforward direction may be disrupted, as both TC and CC inputs to L4 cells – which mediate intracortical feedforward signals – are suppressed by isoflurane. Our data may demonstrate physiological correlates of components of the predictive coding model and provide insight into the disruption of sensory processing during anaesthesia.

### Future Directions

In these experiments, we investigate the effect of isoflurane on discrete components of a higher-order cortical circuit. Investigating the pathway-, cell type-, and layer-specific effects of isoflurane is imperative for understanding the larger systems in which these components operate; our findings will inform future investigations of sensory processing and changes in consciousness during anaesthesia. Although here we activated inputs to layer 1 independently, future experiments *in vivo* will provide insight into how overlapping inputs to layer 1 affect arousal and sensory processing. It is reasonable to expect that the combined effects of isoflurane on TC and CC inputs, for example, may engender supralinear effects on underlying circuit components, synergistically contributing to disruption of sensory processing. Moreover, *in vivo* investigations, such as those that involve artificially manipulating activity in different cell types during anaesthesia, may allow for causal, detailed descriptions of the pathway- and cell type-specific components that are necessary and/or sufficient for consciousness. Lastly, a recent study by Suzuki and Larkum (2020) suggests that anaesthetics may prevent integration of feedback signals by targeting signal amplification in apical dendrites of pyramidal cells, and implies that this process may be mediated by metabotropic glutamate and cholinergic receptors^57^. Future investigations should probe the extent to which distinct inputs to layer 1 engage metabotropic glutamate or cholinergic systems, and how the synaptic effects of anaesthetics on layer 1 contributes to disruption of input-output computations by anaesthetics.

## AUTHORS’ CONTRIBUTIONS

Study design: CAM, MIB

Data collection: CAM

Data analysis: CAM, MIB

Manuscript preparation: CAM, MIB

## ACKNOWLEDGEMENTS

The authors would like to thank contributions of individuals in the Department of Anesthesiology, School of Medicine and Public Health, University of Wisconsin, Madison, WI, USA, especially Dr. Bryan Krause for insight provided for the statistical analyses conducted in this study, and Sean Grady for technical support.

---

**Supplementary Figure 1.**
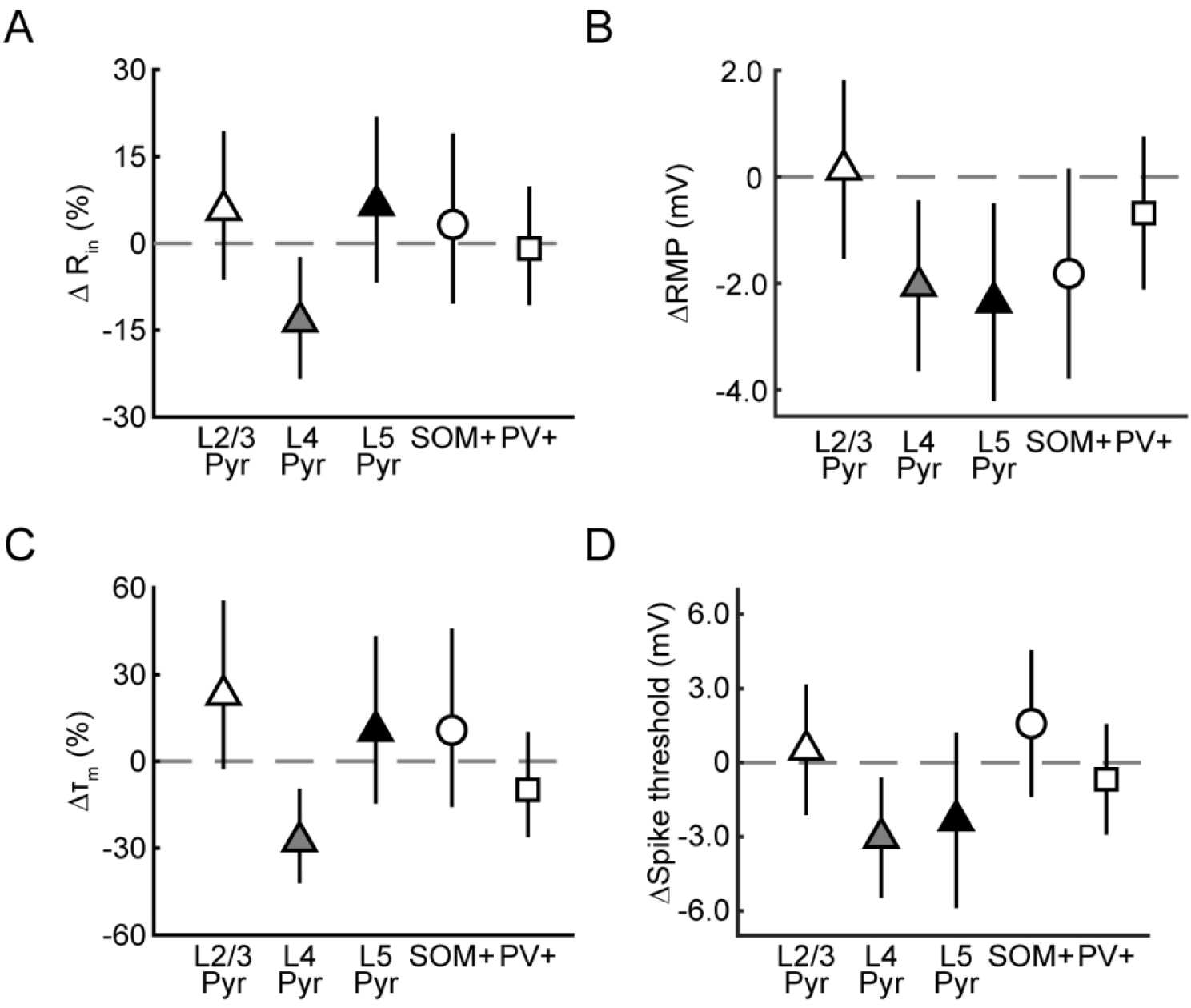
Changes in intrinsic membrane properties of pyramidal cells and interneurons during isoflurane. *A*. Model estimates of percent change of input resistance during isoflurane, where negative values represent decreases in input resistance (i.e., increased conductance) relative to baseline. *B*. Model estimates of shifts of resting membrane potential during isoflurane, where negative values represent hyperpolarization of membrane potentials during isoflurane relative to baseline. *C*. Same as in *A* for percent changes in membrane time constants, where negative values indicate shorter time constants. *D*. Same as in *B* for spike thresholds. Raw values of intrinsic properties for baseline conditions are shown in **Supplementary Table 1**.

**Supplementary Table 1.**
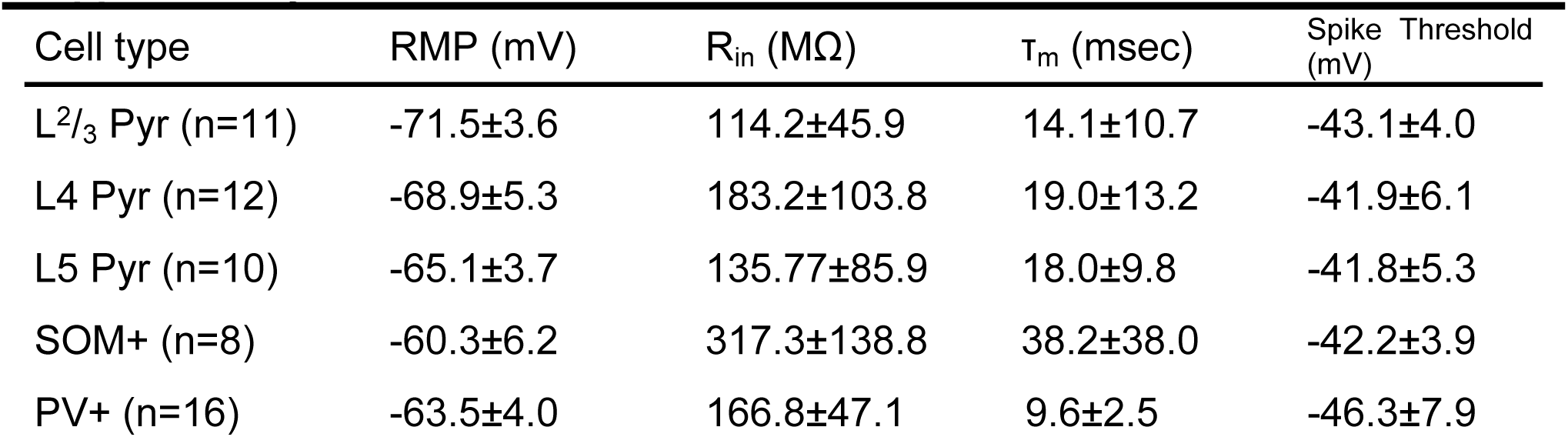
Intrinsic properties of recorded cell types during control conditions. Values are mean ± standard deviation. Properties were measured ∼2 minutes after achieving whole cell access. A series of 10-20 current steps (600 msec) was delivered at progressive intervals (10-40 pA), beginning with a hyperpolarizing current injection. Resting membrane potentials (RMP) were calculated by averaging membrane potentials in the 100 msec prior to onset of all current pulses. Input resistances (R_in_) were extrapolated from relationship between current and steady state voltages. Time constants (τ_m_) were estimated using the first hyperpolarizing current pulse. Spike thresholds were evaluated using the first trial to evoke an action potential, where the spike threshold was the voltage at which the second derivative exceeded twice the standard deviation of the pre-spike baseline value.

| Cell type | RMP (mV) | R<sub>in</sub> (MΩ) | τ<sub>m</sub> (msec) | Spike Threshold (mV) |
| --- | --- | --- | --- | --- |
| L<sup>2/3</sup> Pyr (n=11) | -71.5±3.6 | 114.2±45.9 | 14.1±10.7 | -43.1±4.0 |
| L4 Pyr (n=12) | -68.9±5.3 | 183.2±103.8 | 19.0±13.2 | -41.9±6.1 |
| L5 Pyr (n=10) | -65.1±3.7 | 135.77±85.9 | 18.0±9.8 | -41.8±5.3 |
| SOM+ (n=8) | -60.3±6.2 | 317.3±138.8 | 38.2±38.0 | -42.2±3.9 |
| PV+ (n=16) | -63.5±4.0 | 166.8±47.1 | 9.6±2.5 | -46.3±7.9 |

**Supplementary Table 2.** Changes in short-term plasticity during isoflurane. Estimates are based on predicted values from linear mixed effects model (see Methods) and represent the change in PPR relative to baseline. Negative values represent decreases in PPR during isoflurane.

| Pathway → Cell type | ΔPPR | 95% CI | *P*-value |
| --- | --- | --- | --- |
| TC→ |  |  |  |
| L2/3 Pyr | +0.02 | [-0.32 0.26] | 0.90 |
| L4 Pyr | +0.13 | [-0.10 0.36] | 0.27 |
| L5 Pyr | -0.18 | [-0.43 0.07] | 0.15 |
| PV+ | +0.02 | [-0.18 0.22] | 0.84 |
| CC→ |  |  |  |
| L2/3 Pyr | -0.07 | [-0.30 0.16] | 0.56 |
| L4 Pyr | +0.13 | [-0.10 0.36] | 0.27 |
| L5 Pyr | -0.18 | [-0.42 0.08] | 0.17 |
| SOM+ | +0.45 | [0.22 0.68] | 0.0002 |
| PV+ | -0.03 | [-0.23 0.17] | 0.76 |

